# Novel extracellular, extra-embryonic and intercellular γ-tubulin assemblies in mouse oocytes and preimplantation embryos

**DOI:** 10.64898/2026.10.22.761027

**Authors:** Ruifeng Zhao, Jiaxi Liu, Azizollah Bakhtari, Xiuchun Tian

## Abstract

γ-tubulin is a core component of microtubule-organizing centers (MTOCs), which are traditionally known as intracellular. Here, we report previously unrecognized extracellular, intercellular, and extra-embryonic γ-tubulin assemblies in mouse oocytes and preimplantation embryos in three distinct categories: the γ-tubulin extracellular satellite (γ-TES), γ-tubulin intercellular bridge girdle (γ-IBG), and γ-tubulin nucleation center (γ-TNC). γ-TESs are punctate structures outside the plasma membrane of oocytes and pre-hatching embryos. γ-IBGs are transient, ring-like structures that, together with F-actin, encircle nascent intercellular bridges, suggesting a previously unrecognized role for γ-tubulin in early intercellular bridge formation. γ-TNCs are large, dynamic extracellular assemblies that emerge after fertilization and undergo stage-specific spatial and morphological reorganization during cleavage and blastocyst development. Both γ-TESs and γ-TNCs disappear after embryo hatching. Furthermore, γ-TNCs and γ-IBGs may represent large and ring-like aggregates of γ-TESs, respectively, suggesting that γ-TESs may serve as a local γ-tubulin reservoir for higher order formations. γ-IBGs and γ-TNCs also exhibit close spatial association with microtubules, which form a peri-embryonic web lining the edge of the embryo exterior from zygotes to morulae. Together, our findings reveal a non-canonical, extra-embryonic γ-tubulin organization system that may coordinate an embryonic skeleton architecture across blastomeres for critical embryo-wide developmental events such as polyspermy prevention, polarity establishment, morula and blastocyst formation, and embryo hatching.

## Introduction

γ-tubulin is a highly conserved member of the tubulin family that plays a central role in microtubule nucleation. It assembles into the γ-tubulin ring complex (γ-TuRC), which is the core of microtubule-organizing centers (MTOCs), including canonical, acentriolar or non-centrosomal MTOCs (*1*). In somatic cells, canonical MTOCs are centrosomes and facilitate mitotic spindle assembly (*2*). In mammalian oocytes and early rodent embryos, however, MTOCs are acentriolar (aMTOCs) and are dispersed throughout the cytoplasm and most do not nucleate astral microtubules as opposed to centrosomes (*3*, *4*). Beyond its role in MTOCs, γ-tubulin also assembles into non-canonical structures such as γ-strings and γ-tubules (*5*), which have been implicated in mitochondrial organization and chromatin-associated signaling (*6*, *7*). Collectively, these structures function as a broader organizational system, referred to as the γ-tubulin meshwork (*5*). Previously described γ-tubulin structures reside and function intracellularly (*8*). During preimplantation development, however, embryonic cells often need to operate collectively and coordinately as a single unit which undergoes critical morphological transitions such as polarity formation, compaction, blastulation and hatching. While actin- and microtubule-dependent mechanisms underlying these events have been widely studied, whether γ-tubulin-containing structures contribute to a higher-order organization and coordination at the embryonic level remains unexplored.

Here, we provide the first line of evidence that a whole embryonic skeletal architecture may exist in preimplantation embryos by presenting previously unrecognized extracellular, intercellular, and extra-embryonic γ-tubulin assemblies and a dense microtubule mesh layer lining the embryo exterior. We define these as: (i) γ-tubulin extracellular satellites (γ-TESs), γ-tubulin puncta attached to the plasma membrane of mouse oocytes and pre-hatching embryos underneath the zona pellucida; (ii) γ-tubulin intercellular bridge girdles (γ-IBGs), transient ring-like structures encircling newly established intercellular bridges and associated with the actin cytoskeleton; (iii) γ-tubulin nucleation centers (γ-TNCs), large and dynamic γ-tubulin assemblies that undergo stage-specific spatial reorganization during cleavage and blastocyst development; and (iv) peri-embryonic microtubule web, a dense and tightly interwoven microtubule mesh layer lining the edge of embryo exterior from the zygote to the morula stage. These findings expand the current understanding of cytoskeleton to the extracellular and extra-embryonic space and suggest that an embryonic skeleton with the γ-TNC as the master organizer may link blastomeres into a collective and coordinate embryo-wide morphogenic events during preimplantation development.

## Results

### 1. γ-tubulin extracellular satellites (γ-TESs) are punctate structures outside oocytes and pre-hatching embryos

γ-tubulin structures in oocytes and preimplantation embryos have been widely regarded as aMTOCs (*3*, *4*). However, most aMTOCs outside of the spindle-organizing regions, in fact, do not robustly nucleate microtubules, suggesting two different categories of γ-tubulin assemblies. We performed γ-tubulin immunostaining to comprehensively characterize its dynamics in mouse oocytes and embryos. As reported previously, we observed two distinct γ-tubulin populations, with or without associated microtubule asters. Aster-containing structures were rare, occurring in only 2.3% of GV oocytes and 18.6% of PN zygotes, and were absent in MII oocytes (red arrows in Fig. 1A, Fig. S1A-C, Movie S1). Surprisingly, however, many asterless γ-tubulin puncta “perched” over the plasma membranes of oocytes and embryos and appeared to be extracellular (Fig. 1A-D). To obtain definitive evidence that these γ-tubulin structures were truly extracellular, we co-stained γ-tubulin with E-cadherin, a plasma membrane marker, and revealed that the “cell-surface” γ-tubulin puncta were indeed extracellular and, therefore, also extra-embryonic (Fig. 1E, S1D). This conclusion was further confirmed by that fact the stains were conducted without membrane permeabilization (Fig.S1D), which labels only extracellular proteins. Despite being extracellular, γ-tubulin puncta remained firmly associated with the embryo after zona removal and zona-free culture (Fig.S1E, Movie S2), suggesting the presence of membrane-anchoring or tethering mechanisms. Based on their localization and morphology, we designate these structures as γ-tubulin extracellular satellites (γ-TESs).

**Fig. 1.**
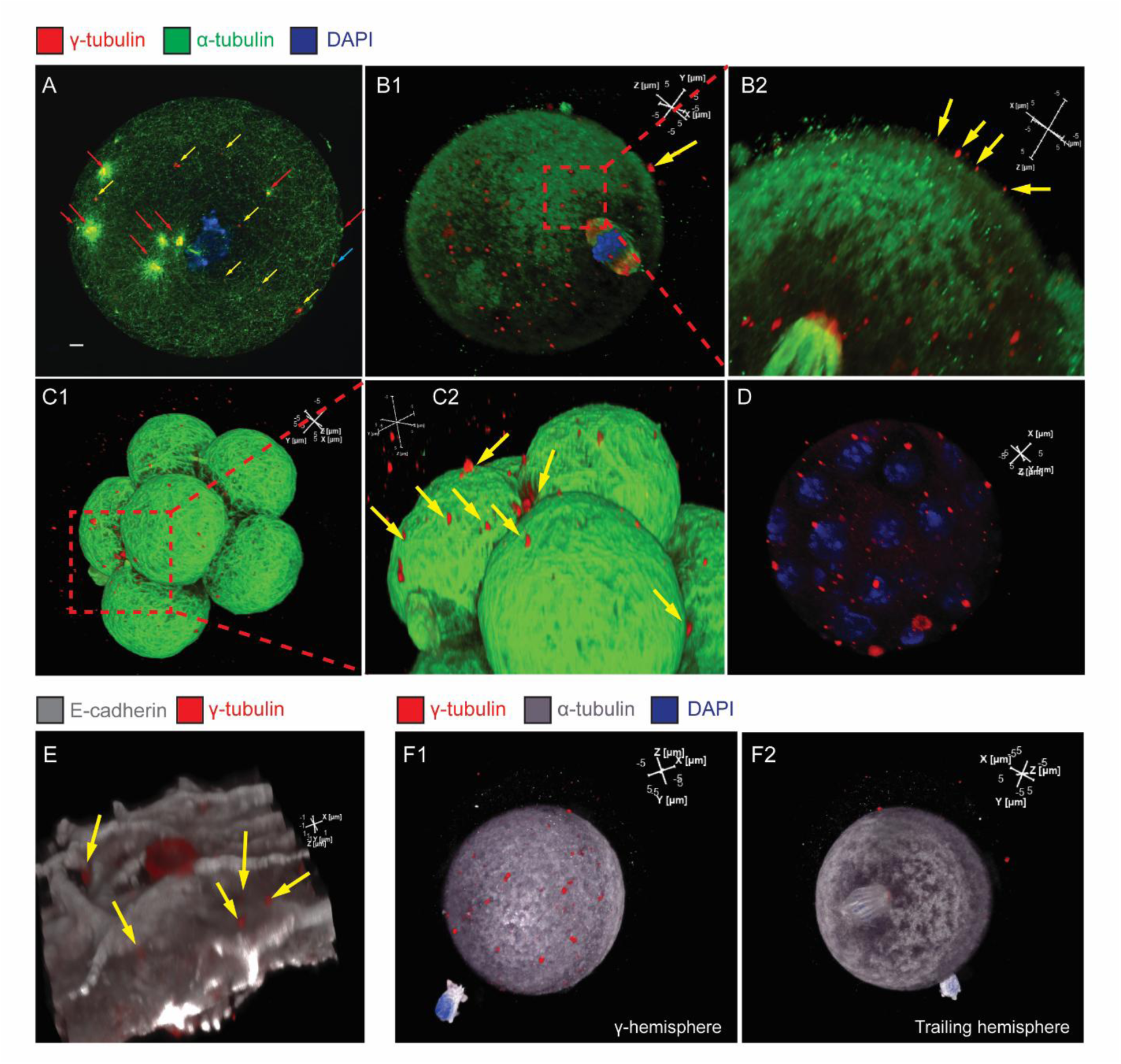
Confocal images of γ-TESs in mouse oocytes and preimplantation embryos. stained for γ-tubulin (red), α-tubulin (green), DNA (blue) and E-cadherin (white) with scale bars=5 μm for each dimension unless indicated otherwise. (A) A single optical section view of a GV oocyte which contained both intracellular aMTOCs with (red arrows) and without microtubule asters (yellow arrows). The blue arrow points to an aster-less and possible extracellular γ-TES. The microtubules (green) were fibrous and distributed throughout the cytoplasm and the nucleus (GV) stained blue in the center. (B) A Z-stacked 3D view of numerous γ-TESs (yellow arrow and red box) appearing to sit on plasma membrane of an MII oocyte lacking visible microtubule asters. The microtubules appeared granular and denser than GV. (C) An 8-cell embryo with the α-tubulin stain (green) illustrating the approximate cell boundaries and the “perched” position of γ-TESs (red box and yellow arrows) over blastomeres. The microtubules were no longer stringy or granular. Rather, they appeared dense and tightly interwoven. (D) A morula showing many “cell surface” γ-TESs. (E) A bird’s eye view of a portion of a blastomere (from a blastocyst) stained for the plasma membrane marker E-cadherin (white). The γ-TESs (yellow arrows) were exterior to the blastomere membrane and are therefore extracellular and extra-embryonic. Scale bar=1 μm. (F) Uneven distribution γ-TESs on the γ- (F1) and the trailing (F2) hemispheres of an MII oocyte. α-tubulin was pseudo-colored in white.

The numbers of γ-TESs and intracellular aMTOCs were roughly similar in both GV (∼50) and MII (∼40) oocytes (Fig. S1B). γ-TESs also exhibited a striking asymmetric distribution. One side of the oocyte/zygote—the γ-hemisphere—contained a higher density of γ-TESs than the opposite or (trailing hemisphere (*p*<0.05; Fig. 1F, S1F, S1H, Movie S3). Of note, the meiotic spindle in MII oocytes was consistently located in the trailing hemisphere. After fertilization, the number of γ-TESs doubled (*p*<0.05) only on the γ-hemisphere but not on the trailing hemisphere or within the cytoplasm (Fig. S1B, S1F). The size of extracellular γ-tubulin structures also increased to more than 5 mm (Fig. S1G). Interestingly, all γ-TESs disappeared after embryo hatching, even though many intracellular MTOCs remained (Fig. S1I, Movie S4).

Because immunofluorescence staining in oocytes and preimplantation embryos could generate artifacts, we included multiple controls. Staining with secondary antibody only (negative controls) produced only faint background signals (Fig. S1J-K). Zona removal (Fig. S1E, Movie S2) and oocyte/embryo fixation with and without methanol treatment did not alter γ-tubulin staining patterns. More convincingly, γ-tubulin structures displayed strong stage- and spatial-specific dynamics—including complete absence in hatched blastocysts (Fig. S1I, Movie S4)—patterns incompatible with nonspecific staining. Together, these data support the conclusion that the extracellular γ-tubulin assemblies reported here are *bona fide* biological structures.

### 2. γ-tubulin intercellular bridge girdles (γ-IBGs) are transient ring-like structures formed during the early stage of intercellular bridge development

Intercellular bridges, or midbodies, are remnants of mitotic/meiotic spindles that persist through the interphase between sister blastomeres in preimplantation embryos. They function as non-canonical MTOCs and also serve as conduits for cytoplasmic exchange (*9*, *10*). A previous study reported that the formation and stabilization of intercellular bridges were not associated with γ-tubulin (*9*). However, upon examining intercellular bridges of different stages of development, we found that a subset of intercellular bridges exhibited strong γ-tubulin signals which encircled the mid-zones of the bridges’ microtubule bundles (Fig. 2A-D, Movie S5). We refer to these ring-like structures as the γ-tubulin intercellular bridge girdles (γ-IBGs). Although intercellular bridges were frequently observed in cleavage stage embryos, the majority of them were not associated with γ-IBGs. We wondered if the γ-IBGs were transient. To determine the exact timing of γ-IBG formation, we utilized the property that the intercellular bridge also forms between the second polar body and the fertilized egg (*11*), and synchronized embryo development by parthenogenetic activation. With the introduction of SPY555-tubulin (Fig. S2C), we precisely pinpointed the initiation of nascent intercellular bridges to approximately 2 hours after activation, while it was still attached to telophase DNA. Intercellular bridges at this stage consistently exhibited intense γ-tubulin and F-actin staining (Fig. 2A-D). These observations suggest a previously unrecognized role for γ-tubulin in the enforcement of intercellular bridges. Additionally, defused and less intense γ-tubulin patches were observed at the end-zones of the bridge’s microtubule bundles (Fig. 2D3, white arrows).

**Fig. 2.**
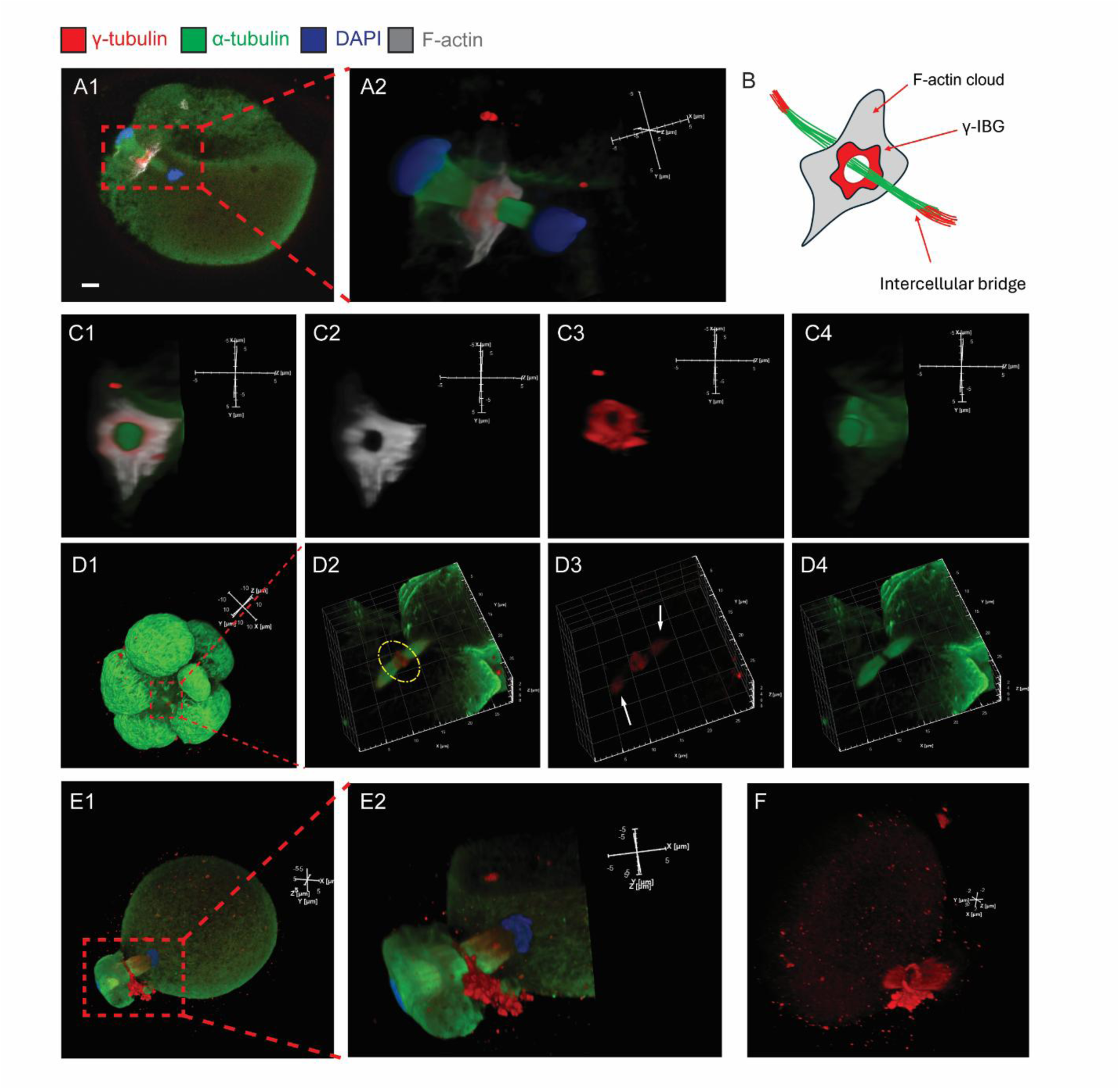
Confocal images of γ-IBGs in mouse pre-implantation embryos. stained for γ-tubulin (red), α-tubulin (green), DNA (blue) and F-actin (white). Scale bars=5 μm for each dimension unless noted otherwise. (A) A single confocal optical section of a newly formed intercellular bridge (A1, red box) between a fertilized zygote and the extruded second polar body, and a fully formed γ-IBGs in the bridge’s mid-zone. The zygote was partially collapsed due to fixation. (A2) An enlarged 3D view of the red box in (A1), illustrating the longitudinal details of microtubule bundles (green) in the intercellular bridge which was still attached to telophase DNA (blue). The γ-IBG (red stained γ-tubulin) and F-actin (white) were in the middle of the microtubule bundles. (B) A schematic representation of the γ-IBG (red), the intercellular bridge (green), the F-actin cloud (white) and the end-zone γ-tubulin patches (red). (C1) A cross-sectional view of the γ-IBG in (A2), elucidating the spatial relationships among the hollowed F-actin cloud (C2), the γ-IBG (red; C3), and the encircled intercellular bridge (green; C4). (D1) An 8-cell embryo with dense interwoven microtubules and the longitudinal view of an intercellular bridge (red box) showing a γ-IBG (red) in the middle. Scale bars=10 μm. (D2) An enlarged view of the red box in (D1), highlighting a yellow circled γ-IBG in mid-zone of the microtubule bundles. (D3) The γ-tubulin (red) view of the structure in (D2) showing the condensed (middle) and patchy red stains at the two ends of the microtubule bundle (white arrows). (D4) The α-tubulin view of the intercellular bridge (green), which appeared broken in the middle, likely due to the presence of the γ-IBG and F-actin which might have obstructed the staining of the encased α-tubulin. (E1) Accumulation of γ-TESs at the future site of γ-IBG formation. (E2) Enlarge view of the red box in (E1), showing details of the aggregated γ-TESs. (F) γ-tubulin stain view of the early formation of γ-IBG, showing the thin γ-tubulin circle, the defused γ-tubulin patches and the aggregates of granular γ-tubulin stains which were likely γ-TESs. Scale bars=2 μm.

While ring-like when fully formed, γ-IBGs appeared as aggregated granules at early stages of formation (Fig. 2E-F, Movie S6). Notably, before γ-IBG’s complete formation, a high number of γ-tubulin puncta, most likely γ-TESs, together with a thin γ-tubulin-rich ring accumulated in the spindle midzone and the future γ-IBG site (Fig. 2E-F). Additionally, the formation of the first γ-IBG between the fertilized zygote and the second polar body coincided with a significant increase in the number of γ-TESs (Fig. 2E-F, S1B). These observations suggest that γ-IBGs likely arose from aggregating γ-TESs.

### 3. The γ-tubulin nucleation centers (γ-TNC) are extracellular and extra-embryonic patches that dynamically change during preimplantation development

γ-tubulin is widely used as a marker to study mitotic and meiotic spindles. However, we found large, dynamic and irregular γ-tubulin patches that, similar to the extracellular and extra-embryonic γ-TESs, also appeared “perched” on blastomere membranes of embryos under the zona pellucida (Fig. 3). Co-staining for γ-tubulin and E-cadherin (Fig. 3N), and γ-tubulin staining without permeabilization (Fig. 3O) confirmed that these structures are also extracellular and extra-embryonic. We designate them as γ-tubulin nucleation centers (γ-TNCs).

**Fig. 3.**
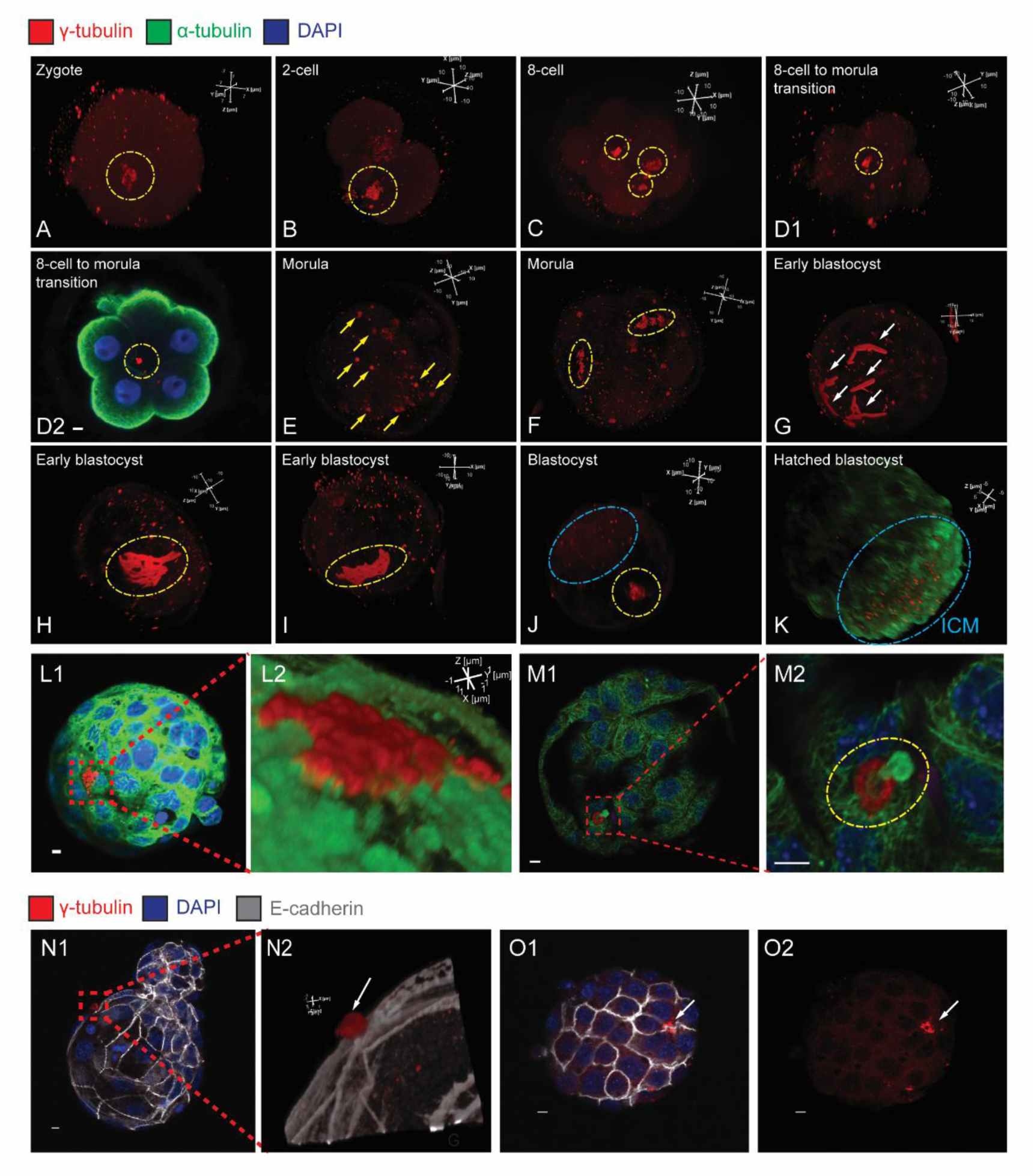
Confocal images of γ-TNCs in mouse pre-implantation embryos. stained for γ-tubulin (red), α-tubulin (green), DNA (blue) and E-cadherin (white). Scale bars=10 μm unless stated otherwise. (A) The Z-stacked 3D view of a PN-stage zygote with abundant γ-TESs and an early and relatively weak-stained γ-TNC (yellow circle). Scale bars=7 μm. (B) A 2-cell embryo with a single and stronger stained γ-TNC (yellow circle) and numerous γ-TESs. (C) An 8-cell embryo with a few patches γ-TNCs (yellow circles). (D1) 8-cell to morula transition: the γ-TNC (yellow circle) was absent from the embryo’s exterior while numerous γ-TESs were visible externally. (D2) A single confocal plane of the embryo in (D1) specifically revealing that the γ-tubulin stain (red) was in the embryo center (yellow circle). The α-tubulin stain (green) was concentrated and lined the “outer edge” of the embryo. The strong α-tubulin stain was missing in regions of cell-cell contact. The nuclei (blue) showed the cells were not dividing and therefore the γ-tubulin stain was not an aMTOC. (E) A 3D-view of a morula which lacked visible external γ-TNCs even numerous large, strongly stained γ-TESs (yellow arrows) were present. (F-G) Morula to early blastocyst transition: the clustering (yellow circles in F) and subsequently zig-zag shaped (white arrows in G) γ-TNCs of stranger and brighter stain indicate their expansion. (H) As the embryo continued to develop, the γ-TNCs merged into one large patch, peaking in stain signal, and possibly size. (I) Subsequently, the γ-TNC patch started to fade and shrink. (J) An expanded blastocyst with a much smaller and weaker-stained γ-TNC (yellow circle) appearing similar to that on the fertilized zygote in (A), when the γ-TNC was just forming. The ICM (blue oval) was located opposite the γ-TNC, which sat over the trophectoderm layer. (K) A hatched blastocyst showing the absence of the γ-TNC. The red stained granules were not external to the embryo and not γ-TESs. Rather, they were intracellular MTOCs confirmed by scanning all single image stacks. (L1) A single optical section of a blastocyst with condensed microtubules and an external γ-TNC. (L2) The magnified view of the red box in (L1) showing the γ-TNC as aggregated granules (likely γ-TESs) which were tightly bound to the dense microtubules inside the cell whose boundary was roughly defined by the enhanced microtubule stain (green). Scale bars=1 μm. (M1) A single optical section of a blastocyst with α-tubulin coming out of the center of a γ-TNC (red box). Scale bars=5 μm. (M2) A magnified 2D view of the red box in (M1) showing details of the key-lock relationship of α-tubulin and the γ-TNC, which was not associated with any dividing cells or spindle. (N1). An early hatching blastocyst with the γ-TNC (red box) sitting outside of the membrane as defined by the E-cadherin stain (white). (N2) A magnified view of the red box in (N1), providing details of the extracellular γ-TNC (white arrow). Scale bars=1 μm. (O1-O2) A blastocyst stained without permeabilization, therefore, only proteins external to the cytoplasmic membrane were stained. The γ-TNC was still visible, confirming it is extracellular.

γ-TNCs emerged after fertilization, persisted to the blastocyst stage, and similar to γ-TESs, disappeared after hatching. Interestingly, they constantly changed in morphology and, at times, localization during preimplantation development. From the zygotic to the 8-cell stage, γ-TNCs were located on top of the plasma membrane and under the zona pellucida (Fig. 3A-C). However, during the 8-cell to morula transition they were surprisingly observed in the embryo interior (Fig. 3D). After morula formation, the γ-TNCs either remarkably returned to or reformed in the sub-zona region (Fig. 3E-F). As embryos transitioned from morulae to blastocysts, γ-TNCs clustered (Fig. 3F), elongated, and adopted zigzag morphologies (Fig. 3G) and were likely in various stages of expansion through clustering. They ultimately formed a large patchy structure (Fig. 3H-I), reminiscent of Jupiter’s Great Red Spot. Also in blastocysts, γ-TNCs consistently localized outside the trophectoderm layer, either opposite or adjacent to the inner cell mass (ICM), but never in the region above the ICM itself (Fig. 3J). After reaching their peaks in size and staining intensity, the γ-TNCs started to shrink (Fig. 3J) and eventually completely disappeared after embryo hatching (Fig. 3K).

γ-TNCs were seen tightly bound to the intracellular microtubule network forming “scab-like” (Fig. 3L) or “key-lock” configurations (Fig. 3M). Similar to γ-IBGs, γ-TNCs also appeared as aggregated granules upon high magnification imaging (Fig. 3L2). γ-TESs puncta were also seen accumulating near γ-TNCs (Fig. S2B; yellow arrows), suggesting that γ-TESs may serve as a local stockpile or reservoir for γ-TNC’s assembly and expansion. While small aggregates of γ-TESs were present in both GV (Fig. S3A; yellow circle) and MII oocytes (Fig. S3B; yellow circle), they were much smaller than the γ-TNCs (>5 mm in diameter; Fig. S1G) in zygotes or cleavage-stage embryos and were not regarded as γ-TNCs.

### 4. The peri-embryonic microtubule web is a dense and interwoven microtubule mesh layer lining the edge of the embryo exterior

The novel γ-tubulin structures and their dynamic changes were associated with striking changes in microtubule organization. In GV oocytes, microtubules appeared as a fibrous and well-defined network throughout the ooplasm (Fig. 4A1, Movie S7), with microtubule asters radiating from some intracellular MTOCs but not from γ-TESs (Fig. 4A2). In mature oocytes, however, this fibrous network was replaced with a punctate or granular staining pattern in the ooplasm which thickened along the cortex (Fig. 4B1, Movie S8). The “volcano crater-like holes” on the cortex (Fig. 4B2) are likely cellular organelles such as cortical endoplasmic reticulum clusters. After fertilization and up to the morula stage, this dense cortical microtubule zone progressively transformed into a thick, tightly interwoven, and almost disorganized mesh layer of fibers lining the edge of the embryo exterior (Fig. 4C-G, Movies S9-10), a structure we termed the peri-embryonic microtubule web. The microtubule mesh layer was absent in regions of cell-cell contact starting from the 2-cell stage (Fig. 4D-G, Movie S11). The initiation and enhancement of this microtubule “fortress” coincided with the increase in the number of γ-TESs (Fig. S1A) and enlargement of γ-TNCs. By early blastocyst, the web was still present in the exterior of the embryo, but microtubule network was seen again in the embryo interior including regions of cell-cell contact (Fig. 4H). This change temporally followed the gradual shrinking of γ-TNCs and reappearance of intracellular MTOCs.

**Fig. 4.**
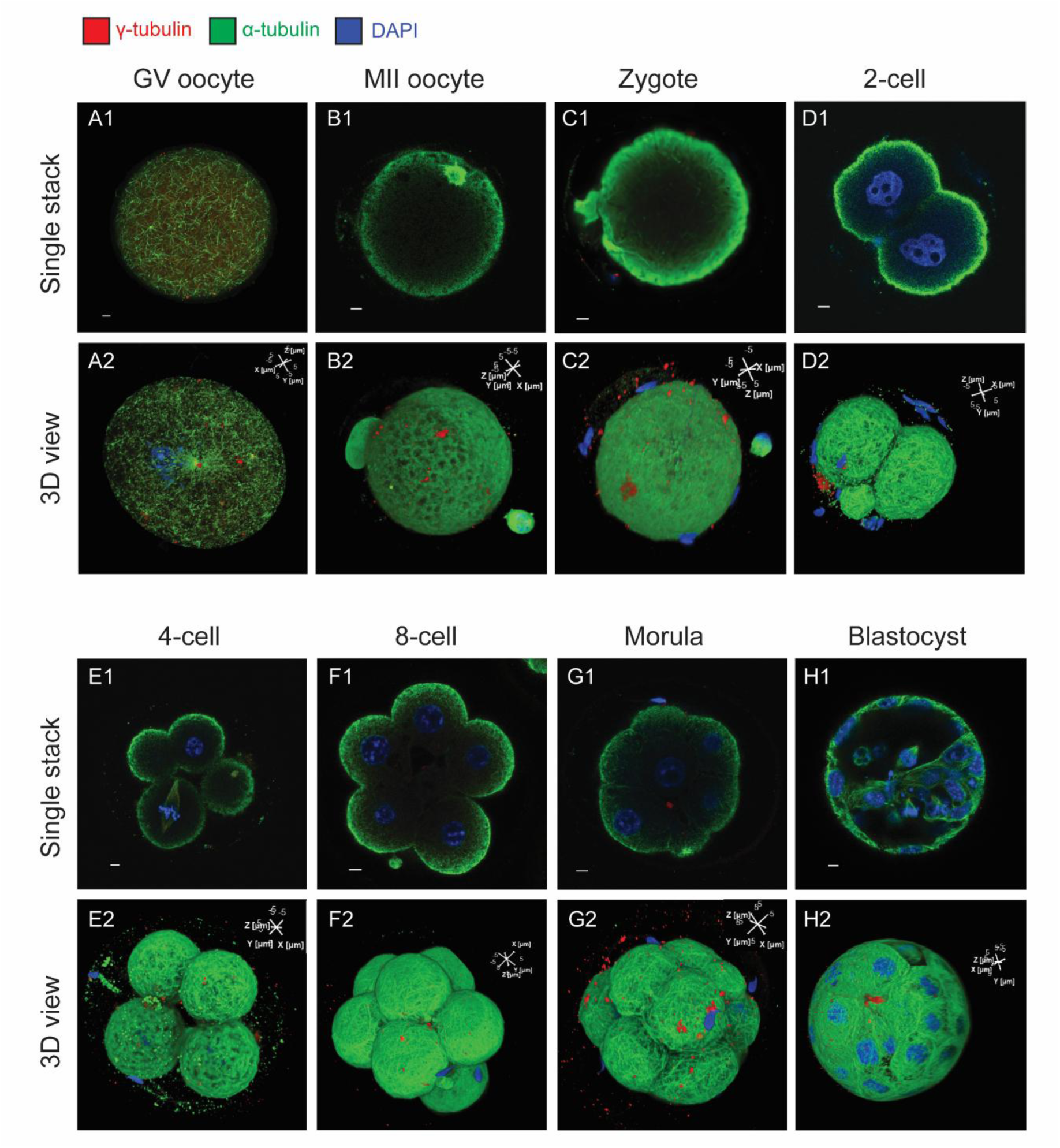
2D (top panels) and 3D (bottom panels) confocal images of the peri-embryonic microtubule web in mouse preimplantation embryos stained for γ-tubulin (red), α-tubulin (green), and DNA (blue). Scale bars=5 μm. (A) In GV oocytes, microtubules were fibrous and either evenly distributed in the cytoplasm or associated with aMTOCs as asters. (B) In MII oocytes, however, the microtubules appeared granular and were concentrated along the cortex. The “volcano-crater” like holes over the granular microtubule zone may indicate locations of organelles such as cortical granules. (C) In zygotes, the granular microtubules started to transform again into either sparse intracellular fibers or dense fibers lining the cortex, the early stage of the peri-embryonic microtubule web. (D-G) As the embryo continued to develop, the microtubule fibers became more obviously interwoven, disorganized and lined the edge of the embryo exterior, progressively forming the mature peri-embryonic microtubule web from the 2-cell to the morula stage. The dense microtubule mesh layer was absent in regions of cell-cell contact. (H) In blastocysts, the peri-embryonic microtubule web was still visible externally but microtubules were also seen intracellularly and along cell-cell contact.

## Materials and Methods

### 1. Collection of GV, in vivo matured and ovulated oocytes, in vitro fertilization (IVF) and embryo culture

The use of CD-1 mice was approved by the Institutional Animal Care and Use Committee (IACUC) at the University of Connecticut. Eight-week-old male CD-1 mice, as well as four to eight-week-old female CD-1 mice, were purchased from Charis Rivers. For GV oocyte collection, female mice were intraperitoneally injected with 7.5 IU pregnant mare serum gonadotropin (PMSG; RP1782721000, BioVendor, Asheville, NC, USA) and euthanized 46 h later. Cumulus– oocyte complexes (COCs) were released by puncturing antral follicles under a dissection scope. COCs containing at least three layers of cumulus cells were selected and incubated in hyaluronidase, and vortexed at the maximum speed for 5 min to remove cumulus cells. The denuded GV oocytes were washed thoroughly in FHM medium (MR-024, Sigma, St. Louis, MO, USA) before use.

All MII oocytes used in this study were in vivo matured and ovulated. Briefly female mice were injected with 7.5 IU PMSG followed by 7.5 IU human chorionic gonadotropin (hCG; C1063, Sigma) 48 h later. Fourteen hours after hCG injection, females were euthanized and COCs were collected from the ampullae of the oviducts into 500 µL IVF medium in a 4-well plate.

For IVF, sperm were released by excising caudal epididymides of euthanized male mice with ophthalmic scissors into 100 μL IVF medium under HypureTM light mineral oil (96005, Kitazato, Fuji, Shizuoka, Japan). The sperm were incubated at 37°C in 5.5% CO₂ and 5% O₂ for 15 minutes, 10 µL of sperm suspension was transferred into a 90 µL droplet for 1 hour in the STM medium for capacitation. For fertilization, 1 million capacitated sperm were introduced into the IVF well containing in vivo matured and ovulated oocytes. Six hours later, the presumptive zygotes were transferred into FHM and removed of cumulus cells by pipetting. The denuded embryos were cultured in KSOM medium (MR-121-D, Sigma) at 37°C in 5.5% CO₂ and 5% O₂ to various developmental stages.

For zona pellucida-free embryo culture, the zona pellucida of presumptive zygotes was removed with acidic Tyrode solution (T1788, Sigma) and cultured in the Well of the Well (WOW) plate (V005035101, VitaVitro, Shenzhen, China) in KSOM at 37°C in 5.5% CO₂ and 5% O₂.

To capture the transition from the 8-cell stage to morula, mouse embryos were monitored using an EVOS M7000 inverted, multi-channel fluorescence and transmitted light imaging system (Thermo Scientific, Ward Hill, MA) for signs of compaction every 2 hours starting from two days post-IVF. Embryos were fixed when approximately 50% had completed compaction.

### 2. Pathogenetic activation, and determination of early-stage intercellular bridge formation in parthenotes

Mouse in vivo MII oocytes were parthenogenetically activated with ethanol treatment. Briefly, COCs were sequentially transferred through multiple droplets of 7% (v/v) ethanol in PBS to minimize dilution. COCs were then incubated in 7% ethanol for 5 min at room temperature. Following activation, oocytes were washed three times in PBS and twice in FHM medium to remove residual ethanol. The activated oocytes were subsequently transferred to M16 medium (MR-016, Sigma) and cultured as described above. The activated oocytes were stained with SPY555-tubulin (CY-SC203, Cytoskeleton, San Diego, CA, USA) and cultured in an on-stage incubator and monitored by live imaging using the EVOS M7000. Timing of the nascent intercellular bridge formation was recorded and used in subsequent experiments with non-SPY555-stained and activated oocytes to avoid potential artifacts from SPY staining.

### 3. Immunofluorescence staining and confocal microscopy

Oocytes and embryos were fixed following a standard centrosome staining protocol. Briefly, samples were fixed in 4% paraformaldehyde (PFA; Cat: J61899, Thermo Scientific) in PBS for 10 min at room temperature. Following fixation, embryos were washed twice in PBST (PBS containing 0.1% Tween-20) and subsequently incubated in ice-cold methanol at −20°C for 10 min. Embryos were permeabilized in 0.5% Triton X-100 (Cat:A416046.AE, Thermo Scientific) in PBS for 20 min at room temperature and then blocked in 10% heat-inactivated goat serum (Cat: S13150, Atalanta Biologicals, Norcross, GA, USA) in 0.5% Triton X-100 for 1 hour. Primary antibodies (Table S1) were diluted in blocking buffer and incubated with oocytes/embryos overnight at 4°C. Subsequently, samples were washed three times (5 min each) in 0.5% Triton X-100 in PBS and incubated with the appropriate secondary antibody (Table S1) for 1 hour at room temperature. Samples were washed again three times in 0.5% Triton X-100 in PBS. As a fixation control, embryos/oocytes were processed using the same protocol but without methanol treatment.

For γ-IBG and E-cadherin staining, embryos were fixed in PFA without methanol treatment. The F-actin was stained with the Alexa Fluor 488 Phalloidin (A12379, Invitrogen, Carlsbad, CA, USA) along with the secondary antibody. For γ-IBG staining, and the interaction between microtubules and γ-TNCs, embryos were mounted on glass slides using ProLong™ Diamond Antifade Mountant with DAPI (P36966, Invitrogen). For γ-TES staining and localization of γ-TNCs at different developmental stages, embryos were transferred to chamber slides containing 1 μg/mL DAPI (62248, Thermo Scientific) in PBS. Slides were kept in the dark at room temperature for 24 hours to allow the mounting medium solidification and subsequently stored at 4°C prior to imaging. Embryos transferred to ibidi µ-Slide 18 Well chamber slides (No81816, ibidi, Grafelfing, Germany) were stored at 4°C. Immunostaining without permeabilization was conducted using the same procedures as described above except that the methanol fixation and permeabilization steps were skipped, and all reagents were omitted from Triton X-100 and Tween-20. Negative controls were conducted using the same procedures previously described without the primary antibodies. The images were taken with Nikon AXR Confocal microscopy and analyzed using NIS-Elements AR 6.01 and Fiji ImageJ. The sizes of γ-TESs in oocytes and zygotes were determined by measuring their longest dimensions using the NIS-Elements AR software.

### 4. Data analysis

For each development stage, at least 15 oocytes/embryos were examined and three replicates were conducted. The numbers and sizes of γ-TESs in oocytes and zygotes were compared using one-way ANOVA followed by Tukey’s multiple comparisons tests. The value of P < 0.05 and P < 0.01 were considered statistically significant (*), and highly significant (**), respectively.

## Discussion

γ-tubulin has been extensively studied as a core component of MTOCs, which are not only needed for mitotic/meiotic spindle formation but are also widely distributed in the cytoplasm and tethered on the Golgi apparatus and inner nuclear membrane through coiled-coil transmembrane molecules (*12*). Non-centrosomal MTOCs are important for intracellular trafficking, cellular morphogenesis and cell migration. For these reasons, γ-tubulin structures have traditionally been regarded as strictly intracellular (*1*, *5*), which has shaped our understanding of cytoskeletal organization across diverse cell types. Yet, during preimplantation development, blastomeres need to operate as a coordinated collective for embryo-wide morphogenic events such as polarity formation, compaction, blastulation and hatching (*13*, *14*). While intra-embryonic, intercellular and intracellular mechanisms have been studied for these embryo-wide events, extra-embryonic organization systems that function across the embryo have not been described. Our discovery of a previously unrecognized γ-tubulin-based dynamic system extends the current understanding of cytoskeleton to the extracellular, intercellular and extra-embryonic spaces. The three structurally and functionally distinct assemblies—γ-TES, γ-IBG, and γ-TNC—that together define a non-canonical, dynamic embryo-wide γ-tubulin network are tightly linked to a web of peri-embryonic microtubules. Together, they may form an embryonic skeleton complex which possibly coordinates embryo-wide morphogenic events. Our findings thus laid a solid foundation for new frontiers in understanding oocyte maturation and preimplantation development.

Among these components, γ-TESs represent the most broadly distributed pool of extracellular γ-tubulin. While prior studies have reported that oocytes/embryos contained γ-tubulin assemblies with and without microtubule asters, all were regarded as intracellular aMTOCs (*15*). However, their structural differences suggest that these “intracellular aMTOCs” are not functionally the same. Our finding that a high percentage of γ-tubulin assemblies without microtubule asters are in fact extracellular clarifies this long-held misclassification. The persistent presence and widespread distribution of γ-TESs throughout oocytes and prehatching development and their enrichment near γ-TNCs and γ-IBGs raise the possibility that γ-TESs serve as the reservoir of γ-tubulin for the formation and enlargement of γ-TNCs and γ-IBGs. The disappearance of γ-TESs after embryo hatching was temporally correlated with the progressive appearance of intracellular MTOCs and centrosomes, suggesting that the extracellular γ-tubulin either returned to the blastomere interior or disintegrated extra-embryonically. Notably, the γ-tubulin molecule does not contain a signal peptide and is meant for intracellular utilization. Which “backdoor” mechanism (*16*) is used to export γ-tubulin outside the plasma membrane and how γ-TESs are tethered on the cell surface remain to be determined.

The γ-IBG represents a transient intercellular structure, specifically associated with the nascent intercellular bridge. Before the full formation of the ring structure, aggregates of γ-TESs accumulated from one side of the microtubule mid-zone, supporting the possibility that the γ-IBG was formed by γ-TESs gathering. The mature γ-IBG was not only associated with microtubule bundles but also co-localized with F-actin, suggesting a collaborative role in intercellular bridge formation. Our observations suggest that γ-tubulin is recruited to the telophase and elongated spindle to prevent it from disassembling by nucleating microtubules from the mid-zone bidirectionally out of different γ-TES units within the γ-IBG, and therefore, maintaining the intercellular bridge beyond cytokinesis. Importantly, γ-IBGs were very transient, acting solely during the initial phase of bridge formation rather than throughout the entire bridge’s life cycle, which may partially explain why intercellular bridges stained mostly negatively for γ-tubulin in a previous study (*9*). The same study also concluded that γ-tubulin was dispensable for intercellular bridge formation by conducting γ-tubulin knockdown experiments in zygotes (*9*). The knockdown, however, may be incomplete due to the presence of an enormous stockpile of maternal γ-tubulin, which allowed the formation of the intercellular bridge despite the knockdown. Also of interest is the diffuse γ-tubulin signals at the end-zones of the intercellular bridge. While the diffuse γ-tubulin signals were frequently observed in mitotic/meiotic spindles, they were always connected to the punctate γ-TuRCs of the centrosomes/aMTOCs. The diffuse γ-tubulin patches in the intercellular bridge, however, were disconnected from the γ-IBG. The role of such γ-tubulin is of interest for future investigation.

The γ-TNC represents the most prominent and dynamic structure within the extracellular and extra-embryonic γ-tubulin network. γ-TNC’s spatial and morphological changes were associated with developmental landmarks, suggesting roles in these processes. First, the two spatial transitions, from embryo exterior to interior and then back to exterior, coincided with the shift of blastomeres from a loosely to cohesively connected state, and the completion of compaction. Second, γ-TNC’s peak size and disappearance were temporally associated with the blastulation transition and hatching process. γ-TNCs’ functional roles are suggested by their close association with the microtubules either in “scab-like” or “key-lock” configurations (Fig. 3L-M). It has been shown that γ-TuRCs tethered to membranes of the Golgi apparatus and the nucleus nucleate microtubules intracellularly. γ-TNCs may also act as membrane-tethered, non-canonical MTOCs that coordinate cytoskeleton organization.

The disappearance of γ-TNC following blastocyst hatching further supports its stage-specific roles before the onset of lineage specification and rapid enlargement of the embryo. After hatching, embryonic cells may no longer need to operate as a single entity and the γ-TNC may also lose its control over the rapidly enlarging embryo. γ-TNC’s disappearance is also in temporal proximity with the progressive emergence and maturation of centrioles (*17*, *18*), which spatially confines to intracellular γ-tubulin (*19–21*), consistent with our observation that dense intracellular microtubule network reappeared in blastocysts (Fig. 4H). It is possible that centriole maturation functionally marks the developmental transition of cytoskeleton organization from embryo-wide and γ-TES/TNC-dependent to one that is blastomere-wide and centrosome-dependent.

The images of microtubules in fibrous networks in GV oocytes and granular/punctate forms in MII oocytes have appeared in prior reports even though the specific term of punctate microtubules was not used (*22–24*). These changes in microtubule morphology may reflect different post-translational modifications of tubulins (*25*). The lining of the embryo exterior by a web of tightly interwoven microtubules after fertilization, however, has not been reported before. With the increase in γ-TES density and γ-TNC size from the zygotic stage, it is highly likely that the peri-embryonic microtubule web is formed from γ-TNCs and γ-TESs nucleating microtubules at multiple extra-embryonic locations. Another observation supporting this possibility is that the microtubule mesh layer was absent in regions of blastomere-blastomere contact. This could result from the lack of γ-TESs and γ-TNCs in these areas. This microtubule web started to develop in zygotes and became extensive as early as the two-cell stage, suggesting a role in the subsequent formation of polarity. More intriguingly, the extra-embryonic γ-TNCs, γ-TESs and the peri-embryonic microtubule web may constitute an embryonic skeleton which organizes the embryo into a single entity for coordinated developmental transitions such as polarity formation, compaction, blastulation and hatching.

Taken together, our findings reveal a previously unrecognized extracellular, extra-embryonic and intercellular γ-tubulin network and peri-embryonic microtubule web that interact and operate intercellularly and at the whole embryo level. In this model, γ-TESs serve as a mobile γ-tubulin stockpile, γ-IBGs direct localized microtubule organization between nascently divided blastomeres, and the γ-TNC functions as a higher-order organizer that links cytoskeletons of individual blastomeres into a single entity for coordinated and embryo-wide morphological changes. It is possible that this network is only present in pre-implantation embryos due to the presence of the zona pellucida for encasement and the need for a multicellular structure to function as a single entity, up to the stage of embryo hatching. Our discoveries challenge the prevailing view that γ-tubulin functions exclusively within the cytoplasm and suggest a hierarchical skeletal architecture and its regulatory roles in oocyte and early embryo development.

## Acknowledgements

We are indebted to Dr. Christopher O’Connell at the Advanced Light Microscopy Facility of UConn COR^2^E for assistance with confocal microscopy and image analysis.

## Funding

USDA W5171 Regional Collaboration Project

USDA-ARS Cooperative Agreements 58-8042-0-028 and 59-8042-5-031.

## Author contributions

Conceptualization: RZ, XT Methodology: RZ, XT

Experimental Execution: RZ, JL, AB Original draft: RZ, XT

Review and editing: RZ, JL, AB, XT Funding acquisition: XT **Competing interests:**

Authors declare that they have no competing interests.

## Data, code, and materials availability

All data are available in the main text or the supplementary materials.

---

**Fig. S1.**
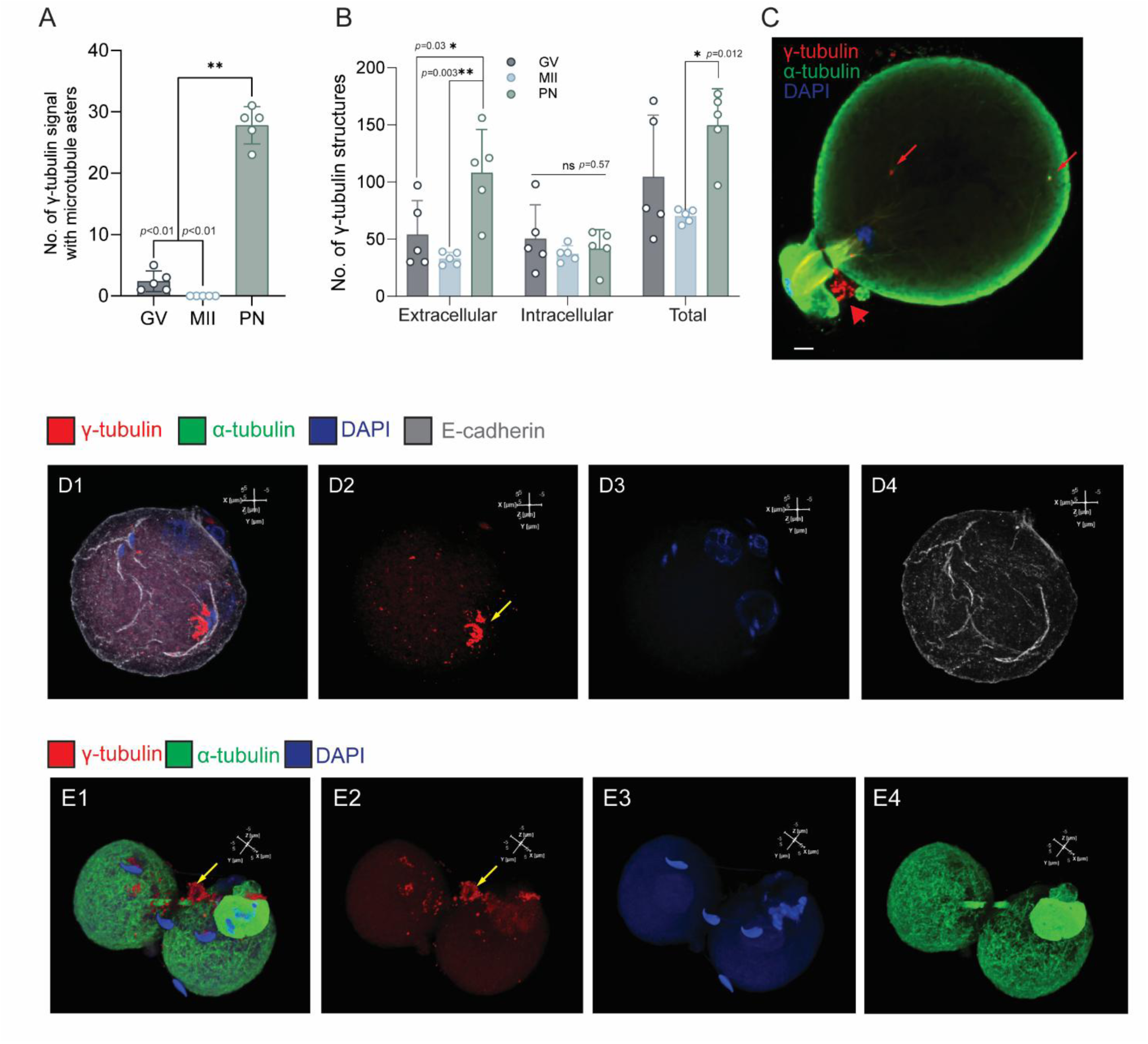
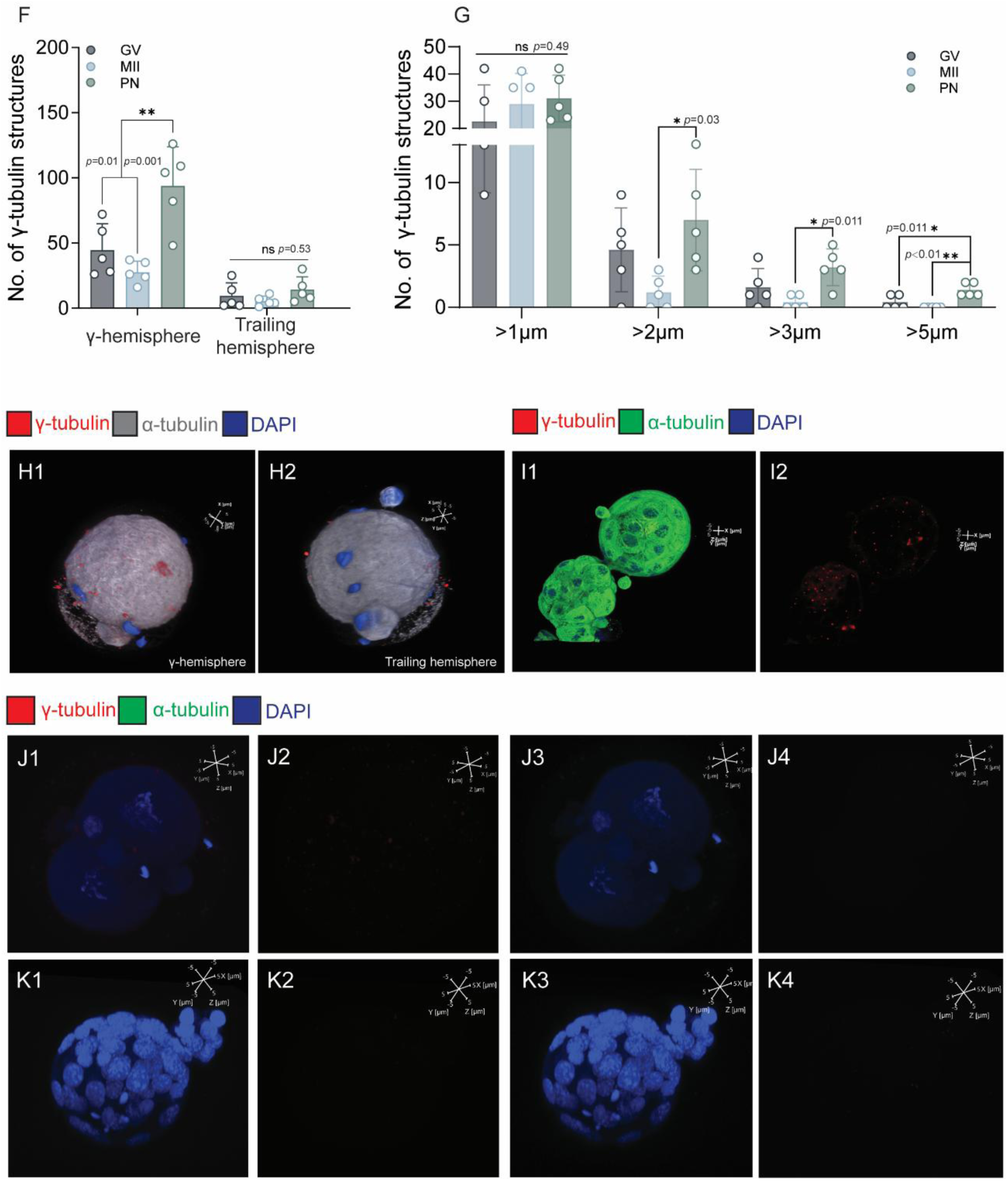
γ-TES and intracellular aMTOC quantifications and confocal images of mouse oocytes and pre-implantation embryos stained for γ-tubulin (red), α-tubulin (green), DNA (blue) and E-cadherin (white) with scale bar=5 μm unless noted otherwise. (A) Numbers of intracellular γ-tubulin structures with microtubule asters. (B) Numbers of γ-tubulin structures (intracellular, with or without microtubule asters) and γ-TESs (extracellular and without asters) in GV, MII and PN zygotes. (C) Microtubule asters re-formed around some γ-tubulin structures (red arrows) in zygotes from asterless γ-tubulin structures in MII oocytes. (D) A PN zygote stained for E-cadherin and γ-tubulin without permeabilization showing that both γ-TESs and γ-TNC (yellow arrow) were outside the plasma membrane and therefore extracellular. (E) A 2-cell embryo with the zona pellucida removed at the zygotic stage did not lose either γ-TESs or γ-TNC (yellow arrow), suggesting membrane tethering mechanisms. (F) Numbers of γ-TESs in the γ- and trailing hemispheres. (G) Numbers and sizes of intra- and extracellular γ-tubulin structures in GV, MII oocytes and PN zygotes. (H) γ-TESs (red) were more abundant in the γ-than the trailing hemisphere in a fertilized zygote. Cell boundaries are represented by α-tubulin pseudo-colored in white. (I) Extracellular and extra-embryonic γ-TESs and TNC disappeared in a hatched blastocyst. All γ-tubulin structures were intracellular. (J) A 2-cell embryo and (K) blastocyst stained without the primary antibodies for γ- and α-tubulins (negative control), showing only minimal background stains and demonstrating that the γ-tubulin structures were not staining artifacts.

**Fig. S2.**
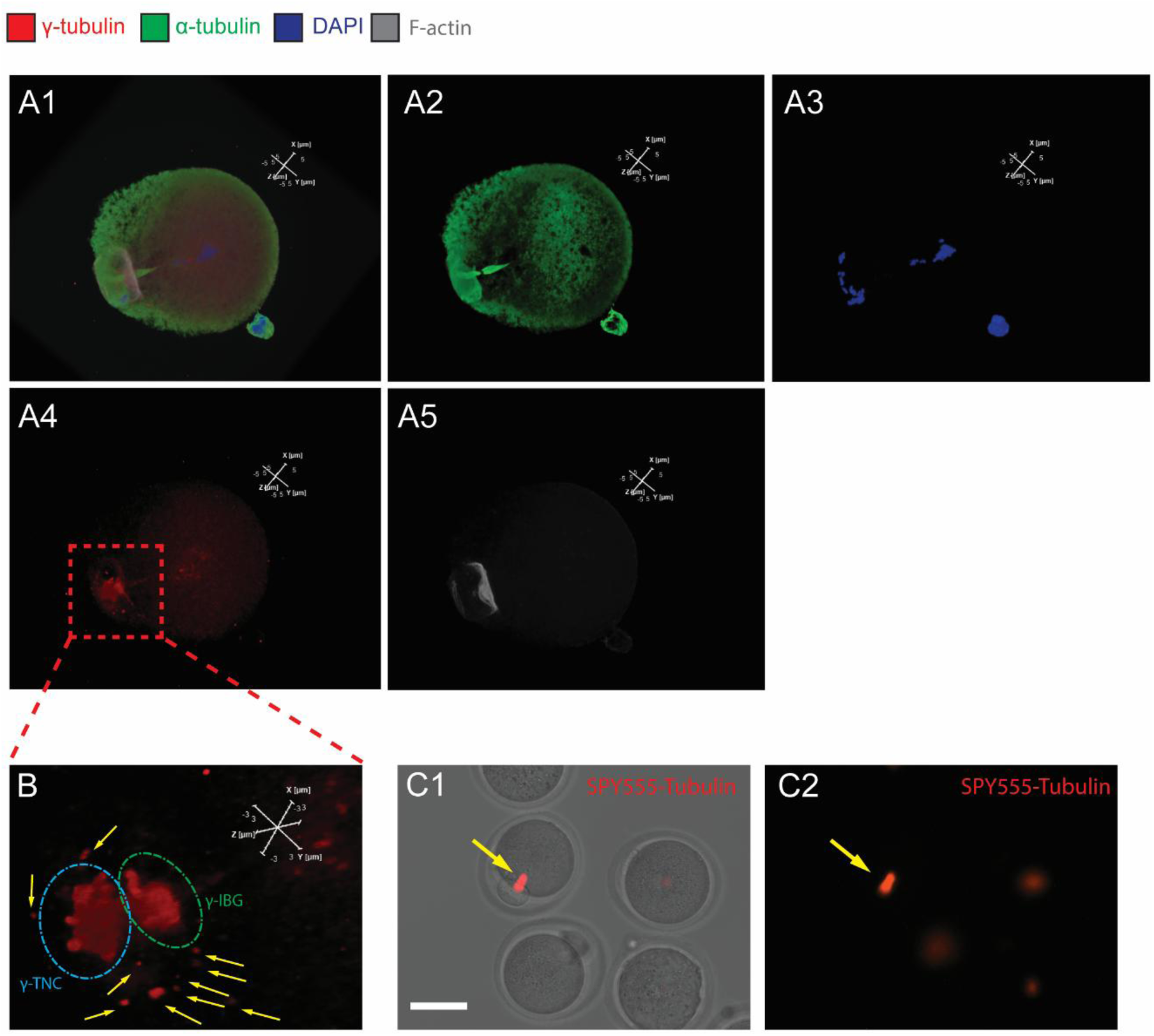
Confocal images of a mouse parthenogenetic zygote stained for γ-tubulin (red), α-tubulin (green), F-actin (white), DNA (blue) or with spindle labeled by SPY555-tubulin (red). Scale bars at 5 μm unless noted otherwise. (A) The zygote had a fully formed γ-IBG between the activated egg and the second polar body. (B) A magnified view of the red box in (A4) showing the presence of both the γ-IBG (green circle) and γ-TNC (blue circle) in close proximity. Both appeared to be composed of granular γ-tubulin structures with multiple γ-TESs (yellow arrows) around, possibly approaching the γ-IBG and γ-TNC. Scale bar=3 μm. (C) A SPY555-tubulin stained pathogenetic zygote, showing the newly formed intercellular bridge. Scale bar=50 μm.

**Fig. S3.**
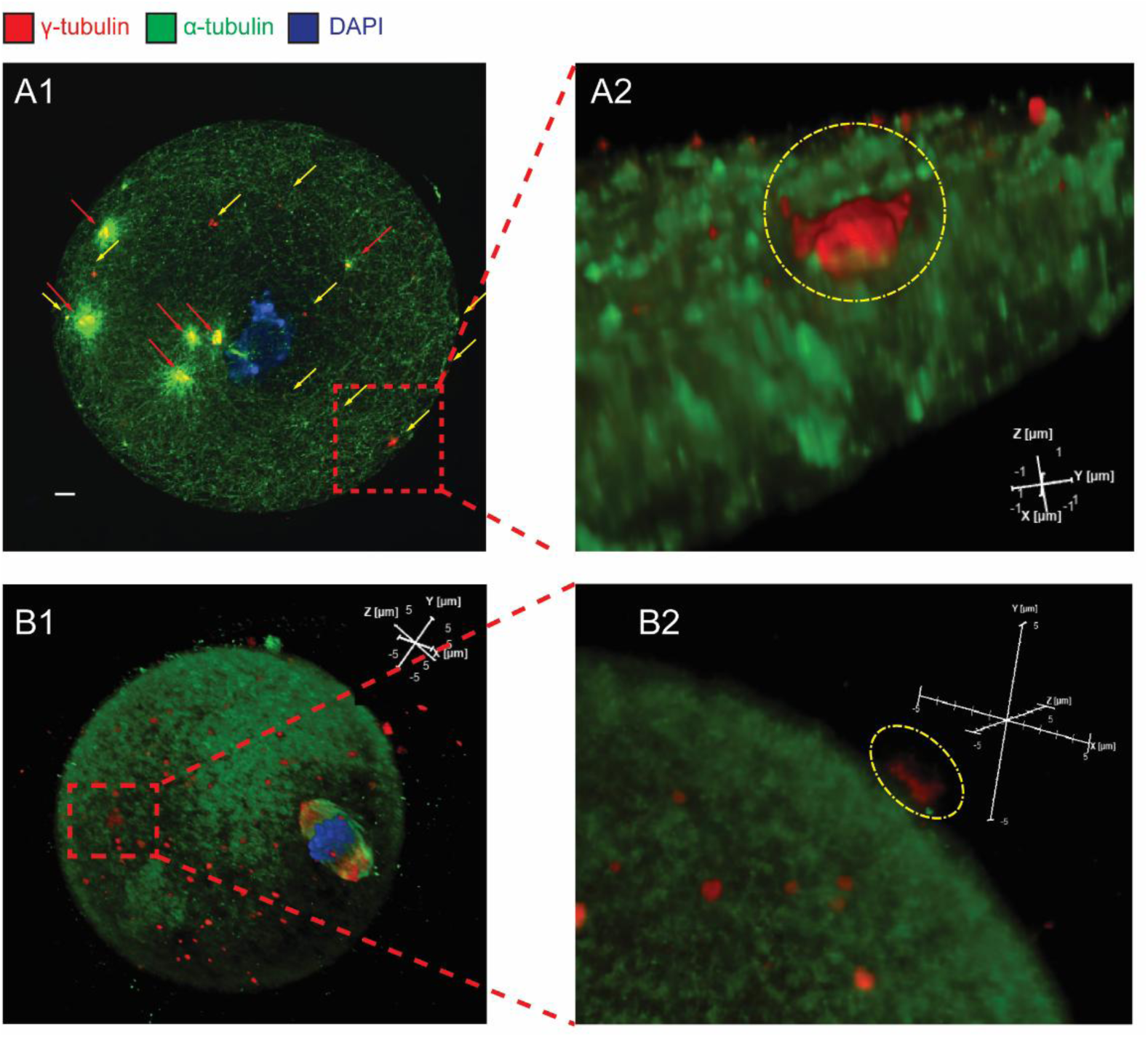
Confocal images of mouse GV (A) and MII (B) oocytes. stained for γ-tubulin (red), α-tubulin (green), and DNA (blue). Both types of oocytes contained small aggregates of γ-TESs (red boxes, scale bar=5 μm; and yellow circles in magnified views, scale bar=1 or 5 μm). They were, however, small and measured less than 5 μm and were not regarded as γ-TNC which were much larger aggregates formed after fertilization.

**Table S1:**
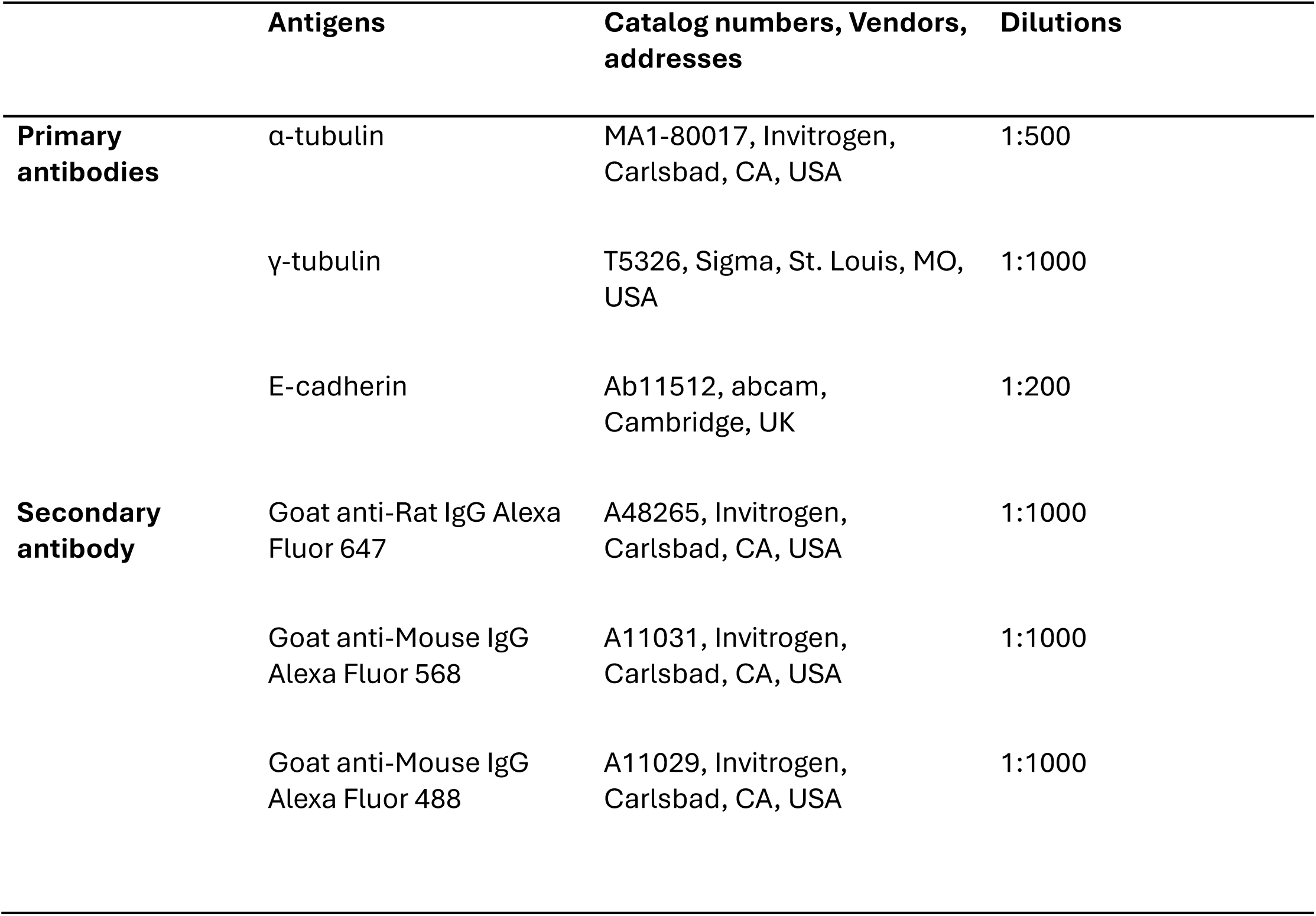
List of Antibodies Used in Immunofluorescence Experiments.

|  | **Antigens** | **Catalog numbers, Vendors, addresses** | **Dilutions** |
| --- | --- | --- | --- |
| **Primary antibodies** | α-tubulin | MA1-80017, Invitrogen, Carlsbad, CA, USA | 1:500 |
| | γ-tubulin | T5326, Sigma, St. Louis, MO, USA | 1:1000 |
|  | E-cadherin | Ab11512, abcam, Cambridge, UK | 1:200 |
| **Secondary antibody** | Goat anti-Rat IgG Alexa Fluor 647 | A48265, Invitrogen, Carlsbad, CA, USA | 1:1000 |
|  | Goat anti-Mouse IgG Alexa Fluor 568 | A11031, Invitrogen, Carlsbad, CA, USA | 1:1000 |
|  | Goat anti-Mouse IgG Alexa Fluor 488 | A11029, Invitrogen, Carlsbad, CA, USA | 1:1000 |

**Movie S1: γ-tubulin with microtubule asters returned at the zygotic stage.**

**Movie S2: γ-TESs (and γ-TNC) were not disturbed by zona removal and zona-free culture in a 2-cell embryo.**

**Movie S3: γ-TESs were unevenly distributed in the γ- and trailing hemispheres of an MII oocyte.**

**Movie S4: A hatched blastocyst contained no γ-TESs. All γ-tubulin structures (red) were intracellular.**

**Movie S5: A 3D view of the fully formed γ-IBG in a zygotic stage embryo.**

**Movie S6: A newly forming γ-IBG in a zygotic stage embryo showing accumulated γ-tubulin granules likely γ-TESs.**

**Movie S7: Fibrous microtubules and γ-tubulin structures with microtubule asters in a GV oocyte.**

**Movie S8: Granular and cortical morphology of microtubules (with “volcano-crater” like holes) in an MII oocyte.**

**Movie S9: The developing per-embryonic microtubule web in a PN zygote.**

**Movie S10: Fully formed per-embryonic microtubule web in a morula. The “emptiness” of one side of the embryo was a result of laser power deficiency in reaching to the entire depth of the embryo.**

**Movie S11: The peri-embryonic microtubule web was absent in regions of cell-cell contact as shown in a 2-cell embryo.**

## References

1. Kollman JM, Merdes A, Mourey L, Agard DA. Microtubule nucleation by γ-tubulin complexes. Nature Reviews Molecular Cell Biology. 2011. pp. 709–721. doi:10.1038/nrm3209

2. Nigg EA, Holland AJ. Once and only once: Mechanisms of centriole duplication and their deregulation in diseases. Nature Reviews Molecular Cell Biology. Nature Publishing Group; 2018. pp. 297–312. doi:10.1038/nrm.2017.127

3. Schuh M, Ellenberg J. Self-Organization of MTOCs Replaces Centrosome Function during Acentrosomal Spindle Assembly in Live Mouse Oocytes. Cell. 2007;130: 484–498. doi:10.1016/j.cell.2007.06.025

4. Gueth-Hallonet C, Antony C, Aghion J, Santa-Maria A, Lajoie-Mazenc I, Wright M, et al. γ-Tubulin is present in acentriolar MTOCs during early mouse development. J Cell Sci. 1993;105: 157–166.

5. Rosselló CA, Lindström L, Eklund G, Corvaisier M, Kristensson MA. γ-Tubulin–γ-Tubulin Interactions as the Basis for the Formation of a Meshwork. International Journal of Molecular Sciences. MDPI AG; 2018. doi:10.3390/ijms19103245

6. Corvaisier M, Zhou J, Malycheva D, Cornella N, Chioureas D, Gustafsson NMS, et al. The γ-tubulin meshwork assists in the recruitment of PCNA to chromatin in mammalian cells. Commun Biol. 2021;4. doi:10.1038/s42003-021-02280-1

7. Lindström L, Li T, Malycheva D, Kancharla A, Nilsson H, Vishnu N, et al. The GTPase domain of gamma-tubulin is required for normal mitochondrial function and spatial organization. Commun Biol. 2018;1. doi:10.1038/s42003-018-0037-3

8. Sulimenko V, Dráberová E, Dráber P. γ-Tubulin in microtubule nucleation and beyond. Frontiers in Cell and Developmental Biology. Frontiers Media S.A.; 2022. doi:10.3389/fcell.2022.880761

9. Zenker J, White MD, Templin RM, Parton RG, Thorn-Seshold O, Bissiere S, et al. A microtubule-organizing center directing intracellular transport in the early mouse embryo. Science (1979). 2017. Available: https://www.science.org

10. Vasilev F, Mihajlović AI, Rémillard-Labrosse G, FitzHarris G. Long-lived cytokinetic bridges coordinate sister-cell elimination in mouse embryos. Dev Cell. 2025;60: 1521–1531.e5. doi:10.1016/j.devcel.2025.01.002

11. Cairo G, Haseeb MA, Schindler K, Lacefield S. A persistent intracellular bridge and cell cycle misregulation enable polar body cell divisions and tumor formation in *Mos* - deficient eggs. BioRxiv. 2025. doi:10.1101/2025.08.26.672293

12. Chumová J, Kourová H, Trögelová L, Halada P, Binarová P. Microtubular and nuclear functions of γ-Tubulin: Are they LINCed? Cells. MDPI; 2019. doi:10.3390/cells8030259

13. Zhu M, Zernicka-Goetz M. Principles of Self-Organization of the Mammalian Embryo. Cell. Cell Press; 2020. pp. 1467–1478. doi:10.1016/j.cell.2020.11.003

14. Płusa B, Piliszek A. Common principles of early mammalian embryo self-organisation. Development (Cambridge). Company of Biologists Ltd; 2020. doi:10.1242/dev.183079

15. Wu T, Dong J, Fu J, Kuang Y, Chen B, Gu H, et al. The mechanism of acentrosomal spindle assembly in human oocytes. Science (1979). 2022;378. doi:10.1126/science.abq7361

16. Cohen M. Through the Back Door: Proteins Escape Cells without Conventional Permission. 2020.

17. Szollosi D, Calarco P, Donahuef RP. ABSENCE OF CENTRIOLES IN THE FIRST AND SECOND MEIOTIC SPINDLES OF MOUSE OOCYTES. J Cell Sci i i. 1972.

18. Howe K, FitzHarris G. A non-canonical mode of microtubule organization operates throughout pre-implantation development in mouse. Cell Cycle. 2013;12: 1616–1624. doi:10.4161/cc.24755

19. Dictenberg JB, Zimmerman W, Sparks CA, Young A, Vidair C, Zheng Y, et al. Pericentrin and-Tubulin Form a Protein Complex and Are Organized into a Novel Lattice at the Centrosome. J Cell Biol. 1998. Available: http://www.jcb.org

20. Palazzo RE, Vogel JM, Schnackenberg BJ, Hull DR, Wu X. Centrosome maturation. Current Topics in Developmental Biology. 1999. doi:10.1016/s0070-2153(99)49021-0

21. Woodruff JB, Ferreira Gomes B, Widlund PO, Mahamid J, Honigmann A, Hyman AA. The Centrosome Is a Selective Condensate that Nucleates Microtubules by Concentrating Tubulin. Cell. 2017;169: 1066–1077.e10. doi:10.1016/j.cell.2017.05.028

22. Li Z, Leng L, Zhai J, Wang X, Yang W, Wang S, et al. Two distinct causes contribute to the low efficiency of human pre-implantation development. Cell. 2026. doi:10.1016/j.cell.2026.05.037

23. Albertini DF. Cytoplasmic microtubular dynamics and chromatin organization during mammalian oogenesis and oocyte maturation. Mutat Res. 1992.

24. Ibáñez E, Sanfins A, Combelles CMH, Overström EW, Albertini DF. Genetic strain variations in the metaphase-II phenotype of mouse oocytes matured in vivo or in vitro. Reproduction. 2005;130: 845–855. doi:10.1530/rep.1.00558

25. F A, Vassena R. Insights of the tubulin code in gametes and embryos: from basic research to potential clinical applications in humans Running title: tubulin in gametes and embryos. Biol Reprod. doi:10.1093/biolre/ioy203/5105747

